# Inhibitory synaptic vesicles have unique dynamics and exocytosis properties

**DOI:** 10.1101/2020.09.21.289314

**Authors:** Chungwon Park, Xingxiang Chen, Chong-Li Tian, Gyu Nam Park, Nicolas Chenouard, Hunki Lee, Xin Yi Yeo, Sangyong Jung, Guoqiang Bi, Richard W. Tsien, Hyokeun Park

**Affiliations:** Division of Life Science, The Hong Kong University of Science and Technology, Clear Water Bay, Kowloon, Hong Kong; School of Life Sciences, University of Science and Technology of China, Hefei, Anhui, China; Chinese Academy of Sciences, Key Laboratory of Brain Function and Disease, University of Science and Technology of China, Hefei, Anhui, China; Department of Chemistry Chung-Ang University, Seoul 06974, Korea; Creative Research Initiative Center for Chemical Dynamics in Living Cells, Chung-Ang University, Seoul 06974, Korea; Univ. Bordeaux, CNRS, Interdisciplinary Institute for Neuroscience, IINS, UMR 5297, F-33000 Bordeaux, France; Institute of Medical Physics and Biophysics, University of Münster, Münster, 48149, Germany; Singapore Bioimaging Consortium, Agency for Science, Technology and Research, 11 Biopolis Way, Singapore; Department of Psychological Medicine, Yong Loo Lin School of Medicine, National University of Singapore, Singapore; Department of Physiology, Yong Loo Lin School of Medicine, National University of Singapore, Singapore; NYU Neuroscience Institute and Department of Physiology and Neuroscience, New York University, New York, NY, USA; Department of Physics, The Hong Kong University of Science and Technology, Clear Water Bay, Kowloon, Hong Kong; State Key Laboratory of Molecular Neuroscience, The Hong Kong University of Science and Technology, Clear Water Bay, Kowloon, Hong Kong

**Keywords:** Inhibitory synaptic vesicles, exocytosis, release probability, three-dimensional tracking, real-time imaging

## Abstract

Maintaining the balance between neuronal excitation and inhibition is essential for proper function of the central nervous system, with inhibitory synaptic transmission playing an important role. Although inhibitory transmission has higher kinetic demands compared to excitatory transmission, its properties are poorly understood. In particular, the dynamics and exocytosis of single inhibitory vesicles have not been investigated, due largely to both technical and practical limitations. Using a combination of quantum dots (QDs) conjugated to antibodies against the luminal domain of the vesicular GABA transporter (VGAT) to selectively label GABAergic (i.e., inhibitory) vesicles together with dual-focus imaging optics, we tracked the real-time three-dimensional position of single inhibitory vesicles up to the moment of exocytosis (i.e., fusion). Using three-dimensional trajectories, we found that inhibitory synaptic vesicles traveled a short distance prior to fusion and had a shorter time to fusion compared to synaptotagmin-1 (Syt1)-labeled vesicles, which were mostly from excitatory neurons. Moreover, our analysis revealed a close correlation between the release probability of inhibitory vesicles and both the proximity to their fusion site and the total travel length. Finally, we found that inhibitory vesicles have a higher prevalence of kiss-and-run fusion compared than Syt1-labeled vesicles. These results indicate that inhibitory synaptic vesicles have a unique set of dynamics and fusion properties to support rapid synaptic inhibition, thereby maintaining a tightly regulated balance between excitation and inhibition in the central nervous system.

**Significance:** Despite playing an important role in maintaining brain function, the dynamics of inhibitory synaptic vesicles are poorly understood. Here, we tracked the three-dimensional position of single inhibitory vesicles up to the moment of exocytosis in real time by loading single inhibitory vesicle with QDs-conjugated to antibodies against the luminal domain of the vesicular GABA transporter (VGAT). We found that inhibitory synaptic vesicles have a smaller total travel length before fusion, a shorter fusion time, and a higher prevalence of kiss-and-run than synaptotagmin-1-lableled vesicles. Our findings provide the first evidence that inhibitory vesicles have a unique set of dynamics and exocytosis properties to support rapid inhibitory synaptic transmission.

## Introduction

Neurons communicate with other neurons and other cell types by releasing neurotransmitters from their presynaptic terminal via the exocytosis (i.e., fusion) of synaptic vesicles at the presynaptic membrane, subsequently activating postsynaptic receptors to mediate downstream effects (1-4). Synapses in the central nervous system can be broadly classified as either excitatory or inhibitory, depending on the type of neurotransmitters that they release and the effects of those neurotransmitters. While excitatory synapses cause the generation, propagation, and potentiation of neuronal responses for processing information (5), inhibitory synapses play an essential role in feedback and feedforward inhibition in order to control neural excitability (6). In the central nervous system, inhibitory synaptic transmission is mediated primarily by release of the neurotransmitter GABA and serves to coordinate the pattern of excitation and the synchronization of the neuronal network, thereby regulating neuronal excitability (7, 8). Thus, maintaining a tightly regulated balance between excitatory and inhibitory neurotransmission is essential for proper brain function.

An extensive analysis of the components and molecular events involved in vesicle fusion and neurotransmitter release has yielded general models describing the organization and functional properties of both presynaptic and postsynaptic components (9-11). On the presynaptic side, a transient increase in local Ca^2+^ concentration due to activation of voltage-gated Ca^2+^ channels triggers the localized buckling of the plasma membrane via a direct interaction between the C2B domain in the protein synaptotagmin-1 (Syt1) and lipids in the membrane (12-14). This leads to the synchronous fusion between the synaptic and plasma membranes and release of the vesicle’s contents into the synaptic cleft (15), where excitatory and inhibitory neurotransmitters act upon postsynaptic glutamate and GABA receptors, respectively.

Importantly, our general understanding of vesicle release stems from studying excitatory neurotransmission and is currently unable to adequately explain the distinct features associated with inhibitory transmission. For example, the size of the readily releasable pool (RRP) of synaptic vesicles in striatal inhibitory GABAergic neurons—probed by a hypertonic sucrose solution—is three times larger than the RRP in excitatory hippocampal glutamatergic neurons (16). Furthermore, inhibitory neurons have both higher average vesicular release probability (P_r_) and more release sites compared to excitatory neurons (16-18). These results suggest that quantitative differences exist between inhibitory and excitatory synapses with respect to vesicle dynamics, release, and recycling. In addition, disproportionate effects between inhibitory and excitatory synapses following knockout of some or all synapsin genes provide a molecular perspective on the putative differences between these two types of synapses (19-22).

Despite the crucial role that inhibitory neurotransmission plays in processing information in the neuronal network, the dynamics of inhibitory synaptic vesicles are poorly understood. In particular, studying the dynamics of single inhibitory synaptic vesicles in hippocampal neurons in real time has been hampered by both technical and practical limitations, including a lack of markers to label individual inhibitory vesicles, the complex three-dimensional structure of inhibitory synapses, and the extremely small size of synaptic vesicles, which have an average diameter of approximately 40– 50 nm, well below the resolution of conventional light microscopy (2).

Here, we report the real-time three-dimensional tracking of single inhibitory synaptic vesicles in cultured hippocampal neurons using dual-focus optics and QD-conjugated antibodies against the luminal domain of the vesicular GABA transporter (VGAT) to selectively label single inhibitory vesicles. We found significant differences between inhibitory synaptic vesicles and Syt1-labeled vesicles with respect to the fusion time, the total travel length prior to fusion, and the prevalence of kiss-and-run fusion. Moreover, we found that the release probability (P_r_) of inhibitory synaptic vesicles was closely correlated with the vesicles’ proximity to their fusion site and with their total length traveled prior to fusion. These findings indicate that inhibitory synaptic vesicles have unique dynamics and fusion properties that support their ability to facilitate rapid neurotransmission.

## Results

### Real-time three-dimensional tracking of single inhibitory synaptic vesicles in cultured hippocampal neurons

To study the dynamics of inhibitory synaptic vesicles in mature synapses, we tracked single inhibitory synaptic vesicles in three dimensions in real time up to the moment of exocytosis in dissociated hippocampal neurons cultured for 14-21 days using our previously reported strategy of loading each synaptic vesicle with a single streptavidin-conjugated quantum dot (QD)-conjugated to biotinylated antibodies against the luminal domain of endogenous synaptotagmin-1 (Syt1) (or Syt1-labeled synaptic vesicles); this approach provides with a spatial accuracy on the order of 20-30 nm, which is less than the diameter of a synaptic vesicle (23). To specifically load the QDs in inhibitory vesicles, we conjugated streptavidin-coated QDs to commercially available biotinylated antibodies against the luminal domain of the vesicular GABA transporter (VGAT), as shown in Fig. 1A. Upon exocytosis induced during electrical stimulation, the QD-conjugated antibodies bind to the luminal domain of VGAT, causing the QD to be taken up into the vesicle upon endocytosis. To confirm selective labeling of inhibitory vesicles, we used a CypHer5E-labeled antibodies against the luminal domain of VGAT (VGAT-CypHer5E) (24) to label spontaneously released inhibitory vesicles, revealing high colocalization between the QD and VGAT-CypHer5E fluorescence signals (Fig. 1B). Only vesicles that were colocalized with both VGAT-CypHer5E were used for analyzing inhibitory vesicles.

**Fig. 1.**
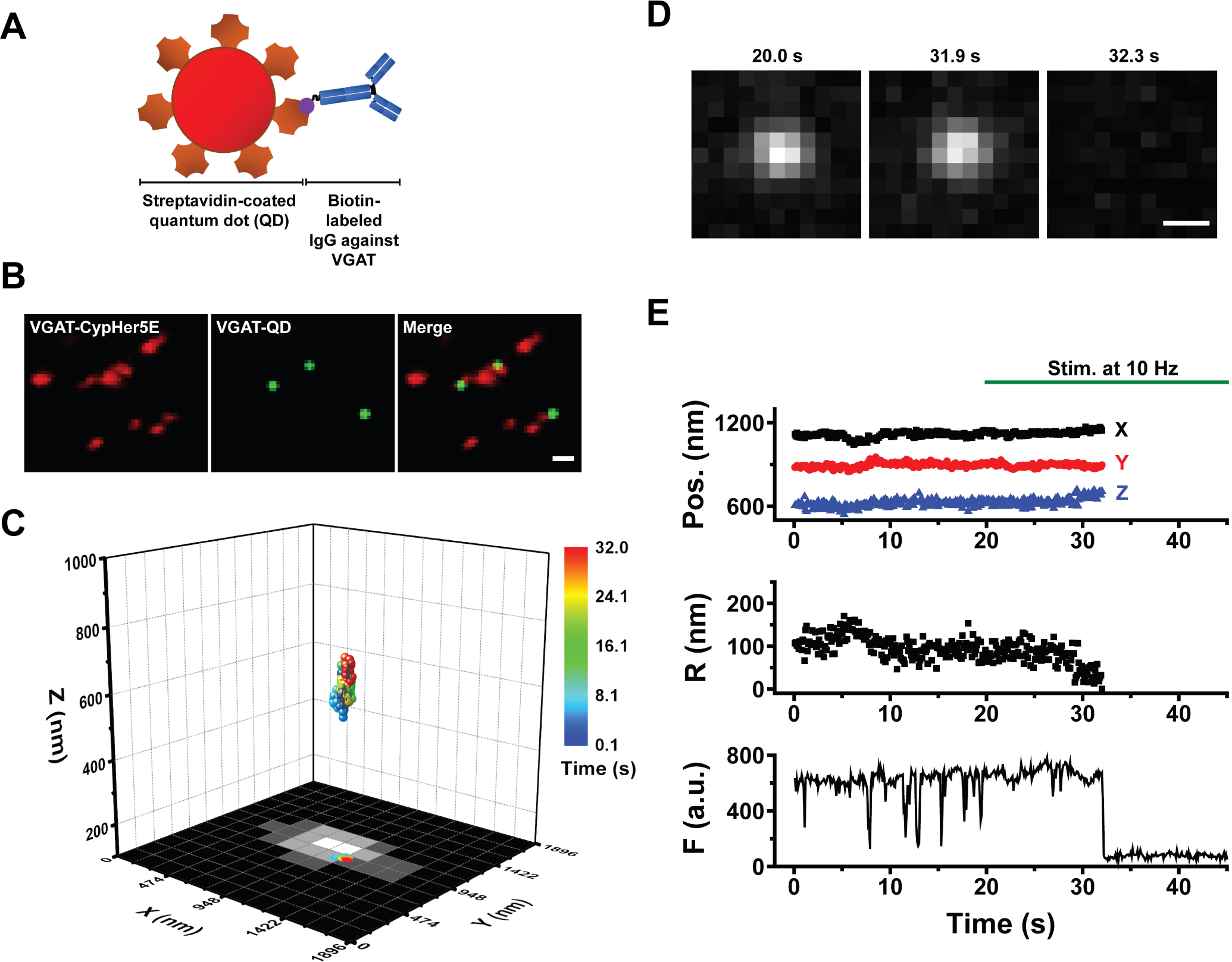
Real-time three-dimensional tracking of a single inhibitory synaptic vesicle loaded with a single quantum dot (QD). **(A)** Schematic diagram depicting a streptavidin-conjugated quantum dot (QD) conjugated to biotinylated antibodies against the luminal domain of VGAT. **(B)** Colocalization of VGAT-QD‒loaded inhibitory vesicles (green) and CypHer5E-VGAT‒labeled presynaptic boutons (red) in cultured hippocampal neurons. Scale bar: 1 µm. **(C)** Three-dimensional trajectory of a VGAT-QD‒loaded inhibitory vesicle overlaid on the *x*-*y* plane of a CypHer5E-VGAT‒ labeled presynaptic bouton. The color bar represents elapsed time; electrical stimulation (10 Hz) started at 20 s, and the vesicle underwent exocytosis at 32.0 s. **(D)** Fluorescence images of the VGAT-QD‒loaded vesicle shown in panel C taken at the indicated times. Scale bar: 0.5 µm. **(E)** Three-dimensional position, radial distance from the momentary position to the fusion site (R), and fluorescence intensity (F) of the VGAT-QD‒loaded vesicle shown in panel C. Note the photoblinking events (e.g., at approximately 8 s, 13 s and 15 s), confirming the presence of one QD inside the vesicle. Electrical stimuli (10 Hz) were applied for 120 s starting at 20 s (green horizontal bar).

In our experiments, exocytosis of a QD-loaded vesicle is reflected by a sudden, irreversible loss of QD fluorescence caused by quenching with extracellular trypan blue (23). To measure the position of single QD-loaded inhibitory synaptic vesicles in the *x*-*y* plane, we used fluorescence imaging with one-nanometer accuracy (FIONA) (23, 25, 26); the vesicle’s *z*-position was localized using dual-focus imaging (23, 27). Figure 1C shows the real-time three-dimensional position of a single QD-loaded inhibitory synaptic vesicle in a cultured neuron stimulated at 10 Hz up until the time of exocytosis; the *x*-*y* plane projection is overlaid on the fluorescence image of an inhibitory presynaptic terminal labeled with VGAT-CypHer5E, showing that the inhibitory vesicle remained near the edge of the presynaptic terminal before undergoing exocytosis. Fluorescence images of the QD-loaded inhibitory vesicle just before and after exocytosis show near-complete, irreversible quenching of the QD 32.0 s after the start of imaging (Fig. 1D). In addition to measuring the three-dimensional position, we also analyzed the average fluorescence intensity (F) within the region of interest (ROI) and the radial distance (R) between the momentary position and the fusion site using the equation 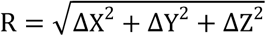 (Fig. 1E). A close examination of the fluorescence trace in Fig. 1E reveals several quantal “photoblinking” events (e.g., at approximately 8, 13, and 15 s), consistent with the presence of a single QD inside the vesicle lumen, followed by a sharp, irreversible loss of fluorescence at 32 s due to exposure of the QD to the quencher (trypan blue) in the external solution, indicating exocytosis. Thus, the example shown in Fig. 1 demonstrates that the three-dimensional position of a single inhibitory vesicle labeled with a single QD can be tracked in real time up to the moment of exocytosis during electrical stimulation.

To determine whether QD-conjugated antibodies against the luminal domain of Syt1 label inhibitory synaptic vesicles, we measured the colocalization between VGAT-CypHer5E-labeled inhibitory presynaptic terminals and synaptic vesicles loaded with QD-conjugated antibodies against Syt1. As shown in Fig. S1, we found a relatively low percentage of colocalization between VGAT-CypHer5E terminals and Syt1-QD‒loaded vesicles (12 ± 1.4 % (N = 13 images)), suggesting that our previous measurements of Syt1-labeled vesicles in the hippocampal neurons likely reflect the dynamics of excitatory vesicles (23). In contrast, we found a significantly higher degree of colocalization (61 ± 3.4% (N = 13)) between VGAT-CypHer5E‒labeled terminals and VGAT-QD‒loaded vesicles (*SI Appendix*, Fig. S1B-C), confirming that VGAT antibody‒conjugated QDs label inhibitory synaptic vesicles preferentially.

Next, we examined the relative proportion of inhibitory neurons in our dissociated hippocampal cultures by co-immunostaining the neurons for the neuronal marker MAP2 and glutamic acid decarboxylase 67 (GAD67), a marker of inhibitory neurons (Fig. 2A). Our analysis revealed 6 ± 0.8% (N = 23) colocalization between MAP2-postive and GAD67-positve puncta (Fig. 2B), suggesting that our cultured hippocampal cultures contain predominantly excitatory neurons. Moreover, we found high colocalization (82 ± 7.7% (N = 19)) between GAD67 and parvalbumin (PV) immunostaining (Fig. 2C), indicating that the majority of inhibitory neurons in our cultures are PV-expressing fast-spiking GABAergic interneurons. By extension, we conclude that our measurements of VGAT-labeled synaptic vesicles are likely to reflect the dynamics of inhibitory synaptic vesicles in fast-spiking PV-expressing GABAergic interneurons.

**Fig. 2.**
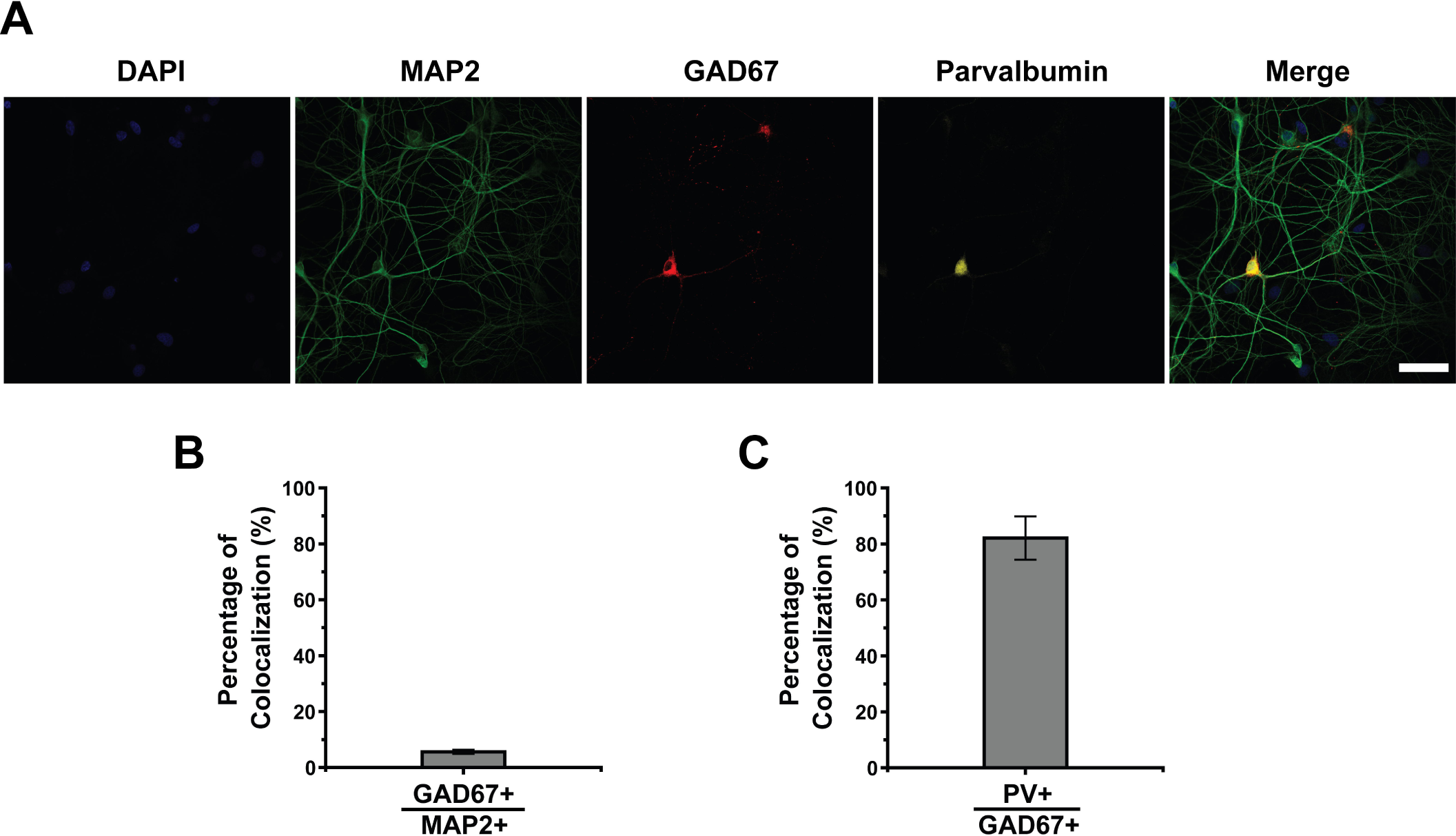
Inhibitory neurons in cultured hippocampal cultures. **(A)** Representative confocal images of cultured hippocampal neurons immunostained for microtubule-associated protein 2 (MAP2, green), glutamic acid decarboxylase 67 (GAD67, red), and parvalbumin (PV, yellow); the nuclei were counterstained with DAPI (blue). Scale bar: 50 µm. **(B**) Percentage of GAD67-positive neurons among all MAP2-positive neurons. **(C**) The percentage of PV-positive neurons among all GAD67-positive neurons.

### Inhibitory synaptic vesicles have unique dynamics

To understand pre-fusion dynamics of inhibitory synaptic vesicles, we measured the net displacement between the initial position and the fusion site of individual vesicles while applying electrical stimuli at 10 Hz. The three-dimensional net displacement of each vesicle was calculated as the Pythagorean displacement using our real-time three-dimensional traces of single QD-loaded inhibitory synaptic vesicles. Figure 3A shows the net displacements of inhibitory synaptic vesicles compared with our previously reported net displacements of Syt1-labeled synaptic vesicles (23). Net displacements between the initial and fusion sites of inhibitory synaptic vesicles were not significantly different from that of Syt1-labeled synaptic vesicles (*p>0*.*7*, Kolmogorov–Smirnov (K-S) test) (Fig. 3A).

**Fig. 3.**
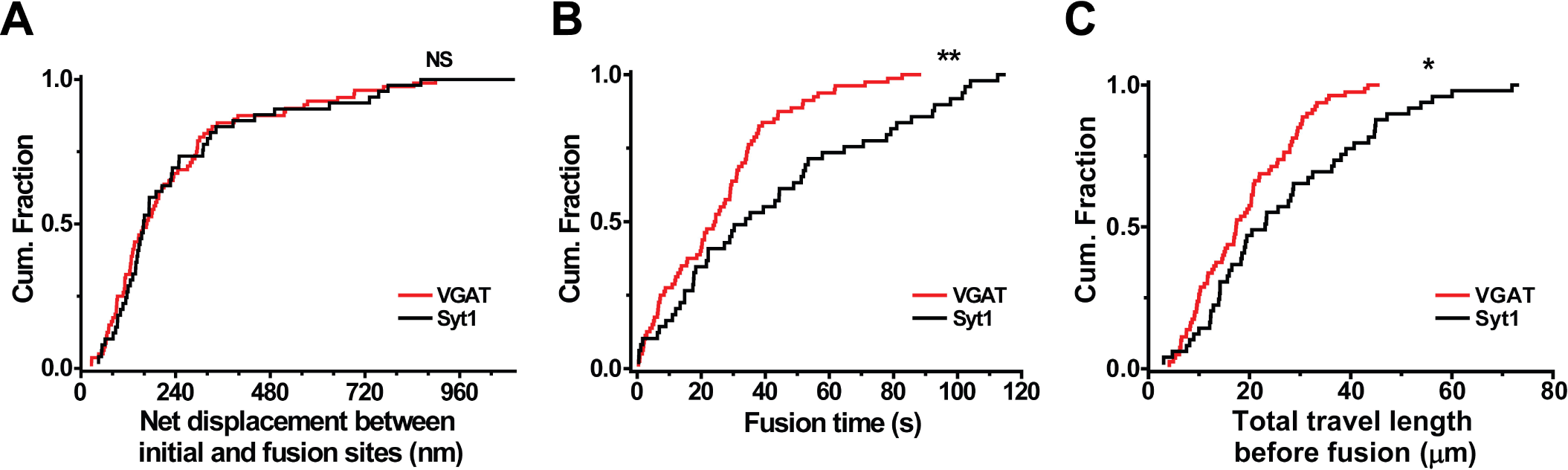
Inhibitory synaptic vesicles have distinct spatiotemporal dynamics. Cumulative distribution of the net displacement between the initial location and fusion site **(A)**, fusion time **(B)**, and total length traveled **(C)** for VGAT-QD‒labeled synaptic vesicles (n = 80 vesicles) and Syt1-QD‒loaded synaptic vesicles (n = 49 vesicles). \**p*<0.05, \*\**p*<0.01, and NS, not significant (Kolmogorov-Smirnov test (K-S test)).

Next, we measured fusion time, which we calculated as the interval between the start of stimulation and the moment of fusion. We found that the average fusion time was significantly shorter for inhibitory vesicles (26.2 ± 2.24 s) compared to Syt1-labeled vesicles (44.4 ± 4.89 s) (*p<0*.*01*, K-S test) (Fig. 3B) indicating that inhibitory vesicles undergo exocytosis more readily (i.e., have higher release probability) than excitatory vesicles. Consistent with this finding, 82.5% of inhibitory vesicles (66 out of 80 vesicles measured) underwent fusion within 40 s of the start of stimulation, compared to 53.1% (26 out of 49) excitatory vesicles, which is consistent with inhibitory vesicles having a higher release probability compared to excitatory vesicles (16, 17).

To understand why inhibitory vesicles fuse more readily than excitatory vesicles yet have similar net displacement prior to fusion, we calculated the total three-dimensional travel distance prior to fusion. We found that inhibitory vesicles traveled a significantly shorter travel length before undergoing exocytosis compared to Syt1-labeled vesicles (19.2 ± 1.13 μm vs. 27.9 ± 2.47 μm, respectively; *p<0*.*05*, K-S test; (Fig. 3C). Taken together, these results indicate that inhibitory vesicles moved their fusion site more straightly compared to excitatory vesicles, thereby increasing their release probability.

### The release probability of inhibitory vesicles is correlated with their position and travel length

Whether a synaptic vesicle’s proximity to its fusion site determines its release probability is a fundamental question in neuroscience. Using real-time tracking of Syt1-labeled vesicles, we previously reported that the vesicle’s proximity to its fusion site is a key factor in determining the release probability (P_r_) (23). However, whether the proximity of inhibitory vesicles to their fusion sites is correlated with P_r_ is currently unknown. We therefore addressed this question and found that net displacement was significantly correlated with fusion time for inhibitory vesicles (Pearson’s r = 0.77) (Fig. 4A). Moreover, the slope of the linear fit was shallower for inhibitory vesicles compared to excitatory vesicles (0.08 ± 0.007 (Fig. 4A) vs. 0.11 ± 0.016 (*SI Appendix*, Fig. S2A), respectively), reflecting earlier exocytosis of inhibitory vesicles. Using net displacement, we then calculated the net velocity of releasing vesicles. We found no significant difference in net velocity between inhibitory and excitatory vesicles (5.4 ± 0.54 nm/s vs. 4.7 ± 0.74 nm/s, respectively) (*p>0*.*2, K-S test*) (Fig. 4B). On the other hand, the coefficient of variation (CV) for net velocity was smaller for inhibitory vesicles (89%) compared to that of excitatory vesicles (110%), indicating that the net velocity of inhibitory vesicles was less variable.

**Fig. 4.**
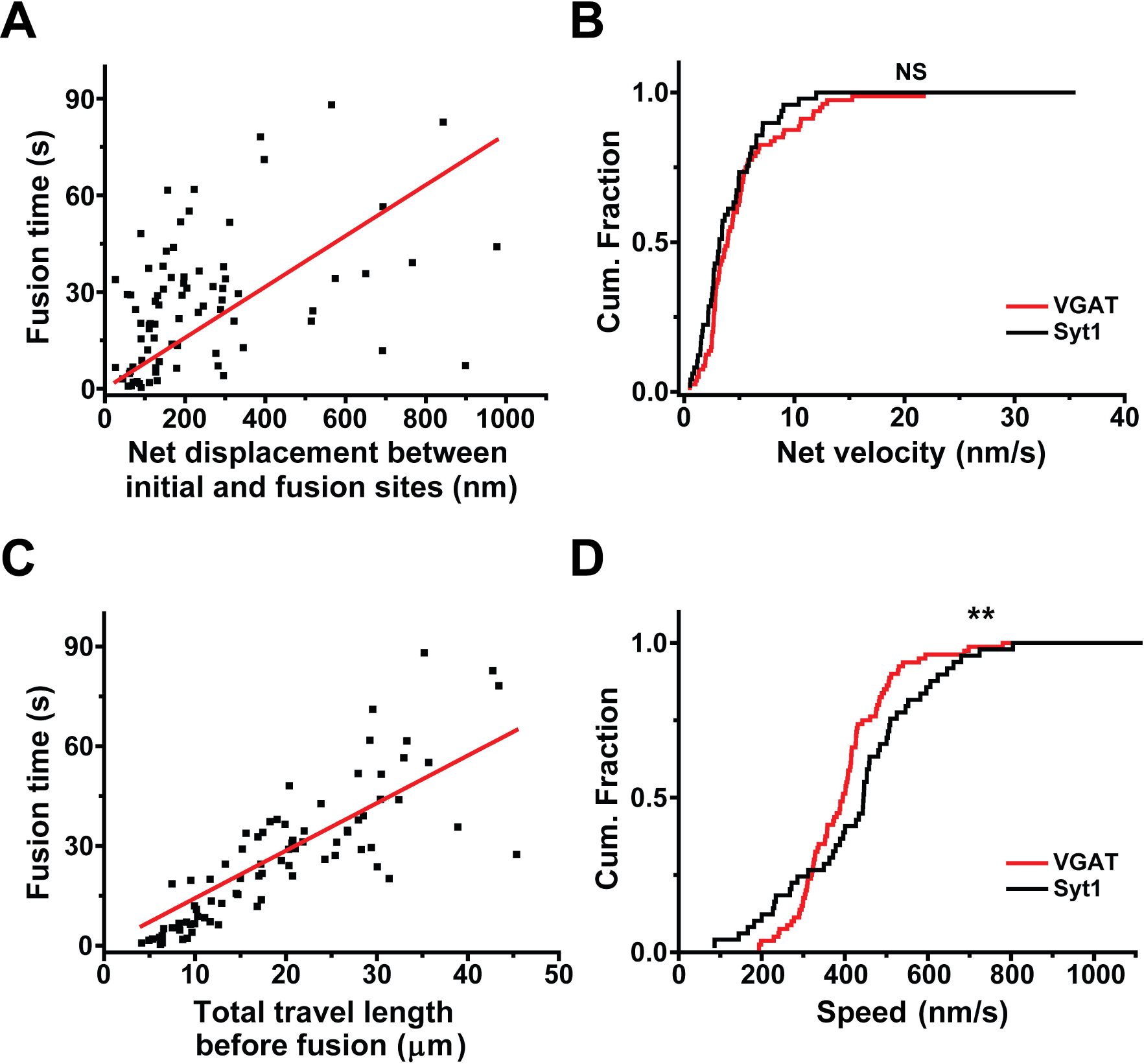
The fusion time of inhibitory synaptic vesicles is correlated with net displacement and total travel length. **(A)** Fusion time is plotted against the net displacement; each symbol represents an individual VGAT-QD‒loaded inhibitory vesicle, and the solid red line represents the linear regression (Pearson’s r = 0.77). **(B)** Cumulative distribution of net velocity measured for VGAT-QD‒labeled synaptic vesicles and Syt1-QD‒loaded synaptic vesicles. **(C)** Fusion time is plotted against the total travel length; each symbol represents an individual VGAT-QD‒loaded inhibitory vesicle, and the solid red line represents the linear regression (Pearson’s r = 0.94). **(D)** Cumulative distribution of vesicle speed measured for VGAT-QD‒labeled synaptic vesicles and Syt1-QD‒loaded synaptic vesicles. \*\**p*<0.01 and NS, not significant (K-S test).

Next, we examined the correlation between fusion time and total travel length prior to fusion and found a significant correlation for inhibitory vesicles (Pearson’s r = 0.94) (Fig. 4C). Consistent with our analysis of net displacement and fusion time, we found that the slope of the linear fit was shallower for inhibitory vesicles compared to excitatory vesicles (1.43 ± 0.06 (Fig. 4C) vs. 1.58 ± 0.09 (*SI Appendix*, Fig. S2B), respectively), indicating earlier exocytosis of inhibitory synaptic vesicles. Using the total travel length prior to fusion, we then calculated the speed (= total travel length/time prior to fusion) of releasing inhibitory synaptic vesicles and found that the speed of inhibitory vesicles were significantly slower compared to excitatory vesicles (405 ± 13.9 nm/s vs. 448 ± 28.1 nm/s, respectively) (*p<0*.*01, K-S test*) (Fig. 4D). As with net velocity, we found that the CV for the speed was smaller for inhibitory vesicles compared to excitatory vesicles (31% vs. 44%, respectively), indicating that the movement of inhibitory vesicles prior to fusion is regulated more tightly compared to excitatory vesicles. Taken together, these results suggest that inhibitory vesicles move toward their fusion site in a more directed fashion than excitatory vesicles.

### Kiss-and-run fusion is more prevalent in inhibitory synaptic vesicles

Finally, we investigated the exocytotic fusion mode of single inhibitory synaptic vesicles. In kiss-and-run fusion, the fusion pore opens briefly, causing a partial drop in fluorescence due to quenching by extracellular trypan blue; in contrast, full-collapse fusion causes the near complete loss of fluorescence (23). Figure 5A shows the fluorescence images of a QD loaded in an inhibitory vesicle undergoing fusion, taken before a drop in fluorescence (70.2 s), after partial drops in fluorescence (71.2 and 105 s), and after the subsequent loss of the remaining fluorescence (105.9 s). The time course of the QD’s fluorescence—obtained by analyzing an ROI containing the QD—shows a sudden drop in fluorescence at 70.5 s (red arrow), followed by a further, irreversible drop in fluorescence at 105.2 s (blue arrow), representing full-collapse fusion (Fig. 5B). The fluorescence of the QD for the period after 70.5 s was reduced to 35% of the initial level, larger than the expected level for QDs steadily exposed to 2 μM TB (12%), which indicates that the inhibitory synaptic vesicle underwent kiss-and-run fusion at 70.5 s.

**Fig. 5.**
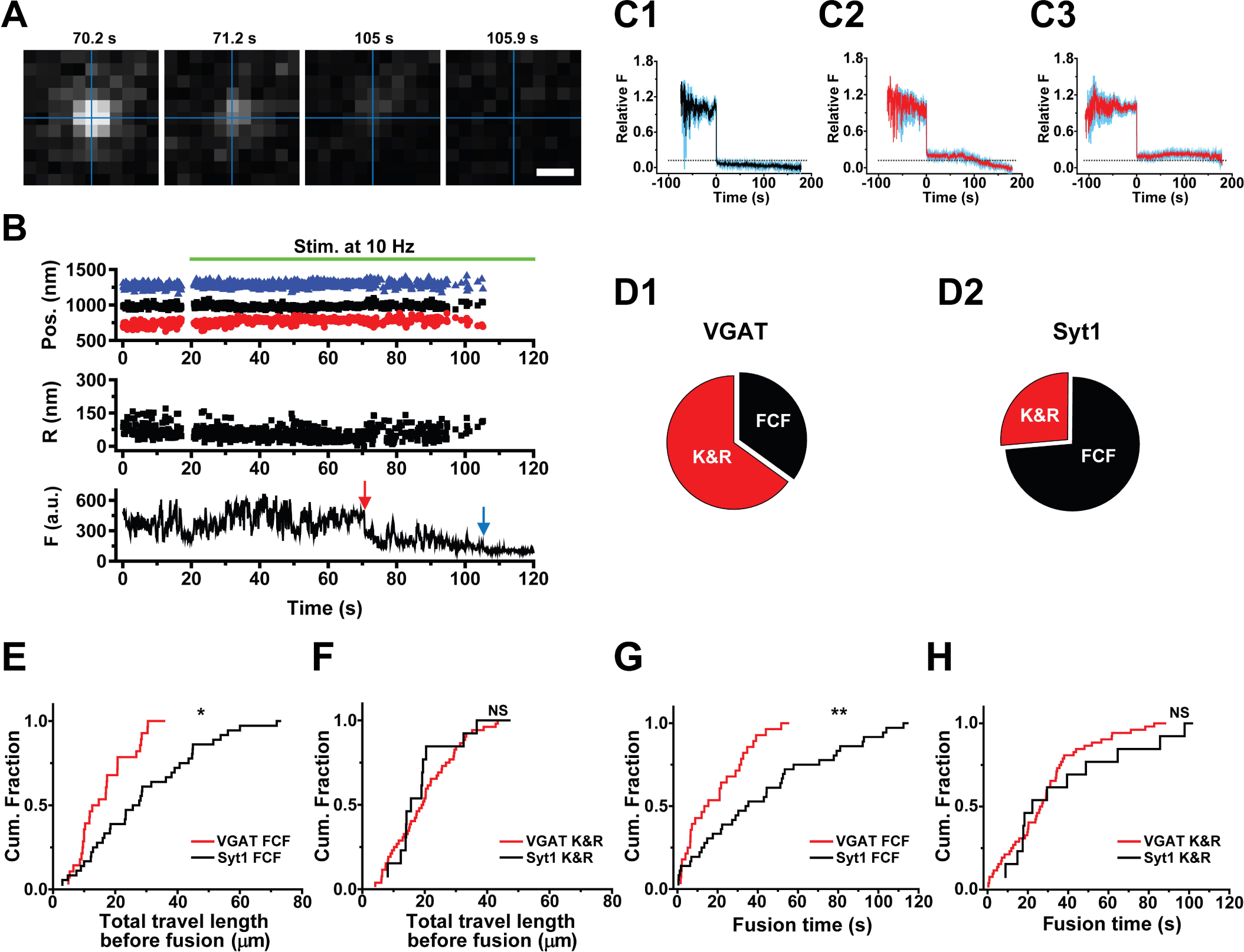
Exocytotic fusion mode of inhibitory synaptic vesicles. **(A)** Fluorescence images taken at the indicated times for an inhibitory vesicle loaded with a VGAT-conjugated QD; electrical stimuli (10 Hz) were applied at 20 s. The intersection of the perpendicular lines marked the position before the first fusion. Scale bar: 0.5 μm. **(B)** Three-dimensional position, radial distance (R), and fluorescence (F) of a VGAT-QD‒ loaded vesicle that underwent kiss-and-run (K&R) fusion (red arrow) followed by full-collapse fusion (FCF, blue arrow). Electrical stimuli (10 Hz) were applied for 120 s starting at 20 s (green horizontal bar). **(C)** Average normalized fluorescence intensity traces (with SEM) time-aligned to the first fusion event for vesicles that underwent full-collapse fusion (FCF) (C1), vesicles that underwent kiss-and-run (K&R) fusion followed by FCF (C2), and vesicles that underwent K&R fusion but never underwent FCF (C3). The dotted horizontal line represents a normalized fluorescence intensity value of 0.12, (the expected normalized fluorescence right after full-collapse fusion). **(D)** Relative distribution of the fusion modes measured in VGAT-QD‒labeled vesicles (D1) and Syt1-QD‒loaded vesicles (D2) that underwent exocytosis, showing that 65% and 27% of vesicles, respectively, underwent K&R fusion. **(E-F)** Cumulative distribution of total travel length for inhibitory and excitatory vesicles that underwent either FCF (E) or K&R fusion (F). **(G-H)** Cumulative distribution of fusion time for inhibitory and excitatory vesicles that underwent either FCF (G) or K&R fusion (H). \**p*<0.05, \*\**p*<0.01, and NS, not significant (K-S test).

We then classified all of the fluorescence traces based on their quenching pattern and aligned the traces relative to the drop in fluorescence, revealing three patterns: 1) near-complete loss of fluorescence (presumably representing full-collapse fusion), 2) partial quenching followed by near-complete quenching (presumably representing a kiss-and-run fusion event followed by full-collapse fusion or reuse after the first kiss-and-run event), and 3) partial quenching that was not followed by near-complete quenching (representing a kiss-and-run fusion event that was not followed by full-collapse fusion); these three patterns are shown in Figure 5C1, C2, and C3, respectively. The average remaining normalized fluorescence following full-collapse fusion was smaller 0.12, suggesting near-complete quenching; in contrast, the remaining fluorescence after kiss-and-run fusion (in both Fig. 5C2 and 5C3) was considerably higher than 0.12. This clear difference in the remaining normalized fluorescence after the first fusion depending on fusion mode confirms that the partial quenching is a good indicator of kiss-and-run fusion in our experiments.

Using the degree of quenching as a measure of full-collapse fusion versus kiss-and-run fusion, we then measured the prevalence of kiss-and-run fusion in releasing inhibitory and excitatory vesicles. We found that approximately two thirds of inhibitory vesicles that underwent exocytosis exhibited kiss-and-run fusion (Fig. 5D1) compared to approximately one quarter of excitatory vesicles (Fig. 5D2), indicating that releasing inhibitory vesicles undergo kiss-and-run fusion more frequently than excitatory vesicles. This relatively high prevalence of kiss-and-run fusion in inhibitory vesicles may contribute to the ability of inhibitory neurons to repeatedly release GABA during prolonged stimulation.

Finally, we examined the dynamics of inhibitory synaptic vesicles based on fusion modes and compared them with excitatory vesicles. We found that among the vesicles that underwent full-collapse fusion, inhibitory vesicles traveled a shorter distance compared to excitatory vesicles (16.9 ± 1.66 μm (n = 28) vs. 30.4 ± 3.09 μm (n = 36), respectively) (*p<*0.05, K-S test) (Fig. 5E). In contrast, when we examined the vesicles that underwent kiss-and-run fusion, we found no significant difference between inhibitory and excitatory vesicles (20.4 ± 1.48 μm (n = 52) vs. 21.0 ± 3.09 μm (n = 13), respectively) (*p>*0.7, K-S test) (Fig. 5F). Similarly, we found that fusion time for inhibitory vesicles undergoing full-collapse fusion was significantly shorter than excitatory vesicles (20.3 ± 3.04 s vs. 44.7 ± 5.84 s, respectively) (*p<*0.01, K-S test) (Fig. 5G), but not for vesicles undergoing kiss-and-run fusion (*p>*0.4, K-S test) (Fig. 5H). On the other hand, the net displacement of inhibitory synaptic vesicles was not significantly different compared excitatory vesicles regardless of the fusion mode of a vesicle (*SI Appendix*, Fig. S4). Taken together, these results indicate that the unique dynamics of inhibitory synaptic vesicles can be attributed primarily to the subgroup of inhibitory vesicles that undergo full-collapse fusion.

## Discussion

GABAergic inhibitory synapses differ from glutamatergic excitatory synapses with respect to both structure and function. In the central nervous system, GABAergic neurons play a critical role in controlling the number and activity of glutamatergic neurons via rapid feedforward and feedback mechanisms (17, 28, 29). To allow for rapid synaptic transmission with high fidelity, inhibitory neurons have a distinct set of properties, including morphology, multiple release sites, biogenesis, and the trafficking of synaptic vesicles in the presynaptic terminal (19, 30-32). Importantly, voltage-gated Ca^2+^ channels are tightly coupled to the release machinery at GABAergic synapses, thereby maximizing the speed and efficacy of GABAergic transmission (33). Although the relatively high release probability and fidelity of action potential‒evoked Ca^2+^ signaling clearly play an important role in maintaining temporally precise GABAergic transmission, the actual exocytosis mode and dynamics of single inhibitory vesicles ranging from their fusion and recycling have not been investigated due to technical limitations.

Here, we selectively loaded inhibitory synaptic vesicles in hippocampal neurons with QDs conjugated to antibodies against the luminal domain of VGAT and then tracked their three-dimensional position in real time up to the moment of exocytosis. We then compared these dynamics with glutamatergic excitatory vesicles labeled with QDs conjugated to antibodies against the luminal domain of Syt1. Although approximately 10% of Syt1-labeled vesicles colocalized with VGAT-CypHer5E-labeled presynaptic terminals, and given that Syt1 serves as the putative Ca^2+^ sensor in both glutamatergic and GABAergic presynaptic terminals (34), our previous results obtained with Syt1-labeled vesicles in cultured hippocampal neurons can be attributed primarily to excitatory vesicles (23). Interestingly, in hippocampal neurons prepared from Syt1 knockout mice, fast synchronous vesicle release was abolished in glutamatergic presynaptic terminals but was spared in GABAergic presynaptic terminals (35). Moreover, a subset of PV-expressing hippocampal interneurons use Syt2 as the primary Ca^2+^ sensor (36), and GABAergic inhibitory neurons can also utilize the high-affinity Ca^2+^ sensor DOC2β (double C2-domain protein beta) to regulate spontaneous release (34, 36). Even though the hippocampus contains more than 21 subtypes of interneurons (7), our immunostaining results indicate that less than 10% of our cultured hippocampal neurons are inhibitory, and approximately 80% of these inhibitory neurons express PV. Thus, our previous data obtained using Syt1-labeled synaptic vesicles in cultured hippocampal neurons likely reflect excitatory vesicles (23). Overall, we found that the total travel length by inhibitory vesicles was smaller compared to excitatory (i.e., Syt1-labeled) vesicles. We also found that inhibitory vesicles underwent exocytosis earlier than excitatory vesicles, with a larger percentage of inhibitory vesicles undergoing release within 40 s of the onset of stimulation. Moreover, we found that the release probability of inhibitory vesicles was closely related to the proximity to their fusion site and their total travel length prior to fusion.

These results indicate that inhibitory synaptic vesicles reach their fusion sites by following a more direct trajectory compared to excitatory vesicles. Interestingly, we also found that kiss-and-run fusion is more prevalent among inhibitory vesicles compared to excitatory vesicles. Taken together, these differences between inhibitory and excitatory vesicles indicate that inhibitory vesicles have a distinct set of dynamics and exocytosis properties, consistent with previous reports that inhibitory synapses have unique electrophysiological and ultrastructural features, including a larger RRP, rapid-onset short-term depression, and higher vesicle release probability (16, 17). Furthermore, these properties support the efficient and rapid release of neurotransmitter from PV-expressing inhibitory neurons, enabling increased microcircuit functions in the brain, including feedforward and feedback inhibition as well as high-frequency network oscillations on a rapid time scale (37). The differences between inhibitory and excitatory vesicles with respect to their release dynamics suggest differences in how these vesicles affect synaptic function, despite having (at least partially) overlapping molecular release machinery. Consistent with the importance of vesicle localization and movement, changes in the relationship between release probability and vesicle position may alter the release of neurotransmitters in neurodegenerative diseases such as Huntington’s disease (38, 39)

The unique dynamics of single inhibitory synaptic vesicles likely contribute to rapid inhibitory neurotransmission. Specifically, their shorter travel length prior to exocytosis and high release probability enable inhibitory vesicles to undergo exocytosis earlier and with higher fidelity compared to excitatory vesicles, features reflected by their significantly shorter time to fusion. In addition, other mechanisms may also contribute to rapid inhibitory neurotransmission. Although the precise mechanism underlying the unique dynamics associated with inhibitory vesicles is currently unknown, several possibilities come to mind, including distinct types and/or states of voltage-gated Ca^2+^ channels, a larger RRP, and tight coupling between Ca^2+^ channels and the Ca^2+^ sensor, features that may support rapid inhibitory neurotransmission and minimize synaptic fatigue at inhibitory presynaptic terminals.

In summary, we provide the first direct evidence that inhibitory vesicles in cultured hippocampal neurons travel less distance to reach their fusion site, undergo exocytosis earlier, and have a higher prevalence of kiss-and-run fusion compared to excitatory vesicles. These unique properties may explain the highly precise manner in which input signals are converted to output signals in inhibitory neurons, providing new insights into how inhibitory neurotransmission regulates the balance between excitation and inhibition in the central nervous system.

## Materials and Methods

### Primary hippocampal neuron culture

Rat (Sprague Dawley) pups at the postnatal 0-day (P0) were sacrificed to harvest CA1 and CA3 of the hippocampal regions of the brain tissue as previously described (23, 40). All procedures were performed according to the animal protocol approved by the Department of Health, Government of Hong Kong. Around 2 × 10^4^ neurons were plated on each Poly D-Lysine (P7886, Sigma-Aldrich, USA)-coated cover glass with a diameter of 12 mm (Glaswarenfabrik Karl Hecht GmbH & Co KG, Germany) in 24-well plates. Two days after plating the neurons, 10 µM uridine (U3003, Sigma-Aldrich) and 10 µM 5-fluorodeoxyuridine (F0503, Sigma-Aldrich) were added to inhibit the proliferation of glial cells in cultured neurons (41). The neurons were cultured in the Neurobasal™-A Medium (10888022, Thermo Fisher Scientific, USA) supplemented with 2.5 % FBS, 1 % Penicillin-Streptomycin (15140122, Thermo Fisher Scientific), 500 µM GlutaMAX™-I Supplement (A1286001, Thermo Fisher Scientific), and 2 % B-27™ Supplement (17504001, Thermo Fisher Scientific). Cultured neurons were maintained in a 5% CO_2_ incubator at 37°C before experiments were performed.

### Microscopy setup

A real-time three-dimensional tracking microscope with a custom-built dual-focus imaging optics was built as previously described (23). The dual-focus imaging optics was made up of an aperture, a beam splitter, lenses, mirrors and a right angle mirror. In the dual-focus imaging optics, the beam splitter divided fluorescent signals into two beam pathways, the focus plane of each beam pathway was determined by its lens, and two fluorescence signals from each beam pathway were imaged side-by-side in an electron multiplying (EM) CCD camera. An Olympus IX-73 inverted fluorescence microscope (Olympus, Japan) equipped with an EMCCD camera (iXon Ultra, Andor, UK) was used to acquire fluorescence signals as previously described (42). An oil immersion 100x objective with NA = 1.40 (UPlanSAPO, Olympus) was used for imaging Streptavidin-coated quantum dots (QD 625, A10196, Life Technologies)-conjugated antibodies against VGAT and CypHer5E-conjugated antibodies against VGAT. In order to locate two-dimensional centroids and to calculate the fluorescent peak intensity at different focal planes (*I*_*1*_ and *I*_*2*_), we used custom-written IDL coded programs (L3Harris Geospatial, USA) to fit a two-dimensional Gaussian function to the fluorescence image. An objective scanner (P-725.2CD, Physik Instrumente (PI), Germany) was used to generate a calibration curve based on the z-position dependence of the normalized intensity difference ((*I*_*1*_*-I*_*2*_*)/(I*_*1*_*+I*_*2*_)). Custom-made programs written in LabVIEW (National Instruments, USA) were used to control an objective scanner and acquire data on the z-position. The z-positions of quantum dots (QDs) were obtained from the relative peak intensity differences using the empirically determined calibration curve, as previously described (23, 27). A 405 nm laser (OBIS, Coherent Inc., USA) and a 640 nm laser (OBIS, Coherent Inc.) were used to illuminate streptavidin coated QDs and CypHer5E, respectively. A 15× beam expander (Edmund Optics) and a focus lens (Newport Corporation, USA) were used to illuminate the sample uniformly.

### Real-time imaging of single inhibitory synaptic vesicles

Imaging experiments were performed using 14 - 21 days in vitro (DIV) rat primary hippocampal neurons at room temperature. Neurons were pre-incubated with 3 nM of CypHer5E-tagged anti-VGAT antibodies (131 103CpH, Synaptic Systems, Germany) in the culture medium and incubated in the 5% CO_2_ incubator for 2 h at 37°C to specifically label inhibitory synaptic vesicles in presynaptic terminals. Then, single inhibitory synaptic vesicles were labeled by QDs conjugated with antibodies against the luminal domain of VGAT. To conjugate QDs to antibodies, Streptavidin-coated quantum dots 625 (A10196, Thermo Fisher Scientific) were incubated with biotinylated antibodies against the luminal domain of VGAT (131 103BT, Synaptic Systems) in a modified Tyrode’s solution (4 mM KCl, 150 mM NaCl, 2 mM CaCl_2_, 2 mM MgCl_2_, 10 mM D-glucose, 10 mM HEPES, 310-315 mOsm/kg, and pH 7.3-7.4) with casein (C7078, Sigma-Aldrich) for 50 min at room temperature. After the incubation, the conjugated solution was added to a sample chamber containing a cover glass with neurons dipped in the modified Tyrode’s solution. The labeling of single inhibitory synaptic vesicles with QDs was conducted by triggering evoked release of synaptic vesicles using 10 Hz electric field stimulation for 120 s, which was induced by parallel platinum electrodes connected with the SD9 Grass stimulator (Grass Instruments, USA). Following 2 min incubation after stimulation, the Tyrode’s solution was superfused for 15 min at 1.0 ∼ 2.0 ml/min. Subsequently, 2 µM of trypan blue (15250061, Thermo Fisher Scientific) were superfused before imaging. During the imaging, trypan blue (quencher) was kept in the chamber to quench the fluorescence of QDs exposed to the external solution during exocytosis. Fluorescence images of CypHer5E-labeled and QD-labeled synaptic vesicles were acquired using an Olympus IX-73 inverted microscope. A ZT640rdc-UF1 (Chroma Technology, USA) and an ET690/50M (Chroma Technology) were used to acquire fluorescence images of CypHer5E-labeled presynaptic boutons. A ZT405rdc-UF1 (Chroma Technology) and an ET605/70M (Chroma Technology) were used to acquire fluorescence images of QD-labeled inhibitory synaptic vesicles. 2000 frame images were obtained using a frame-transfer mode at 10 Hz for 200 s using an EMCCD camera. The EMCCD camera and the stimulator were synchronized with Axon Digidata 1550 (Molecular Devices, USA) to trigger an electric field stimulation at 10 Hz for 120 s while recording. Clampex (program, Molecular Devices) was used to generate stimulation protocols and control stimulation based on the protocol.

### Analysis of the real-time single vesicle-tracking images

Custom-written IDL coded programs were used to localize the centroids of raw fluorescence images. Two-dimensional centroids and peak intensities of synaptic vesicles loaded with QDs at two different focal planes were calculated by fitting a Gaussian function to fluorescence images using custom-written programs using IDL. The position along the z-axis was calculated based on a calibration curve while the average intensity of the fluorescence of a QD within a region of interest (ROI) was calculated using MetaMorph (Molecular Devices). Fusion positions and times of single synaptic vesicles were determined as the first quenching positions and times of fluorescence intensity by trypan blue during electrical stimulation similar to the previous report (23). Two-sample Kolmogorov-Smirnov tests (K-S test) were used for statistical analysis, using OriginPro 9.1 (OriginLab, USA). Data are presented as the mean ± standard error of the mean (SEM). Differences with *p*<0.05 were considered significant.

### Immunofluorescence

For immunocytochemistry, neurons at DIV14 were washed with PBS once, then fixed with ice-cold 100% methanol for 10 minutes at −20°C. After three washes in PBS for 30 min at room temperature, cells were incubated with Chicken antibody against MAP2 (1:1000, ab5392, Abcam), Rabbit antibody against Parvalbumin (1:500, PA1-933, Thermo Fischer Scientific) and Mouse antibody against GAD67 (1:500, MAB5406, Sigma-Aldrich) in staining buffer (0.2% BSA, 0.8 M NaCl, 0.5% Triton X-100, 30 mM phosphate buffer, pH 7.4) overnight at 4°C. Neurons were then washed three times in PBS for 30 min at room temperature and incubated with Alexa Fluor conjugated secondary antibodies (Goat anti-Chicken-Alexa 488, A11039, Life Technologies; Goat anti-Rabbit-Alexa 568, A11011, and Donkey anti-Mouse-Alexa 647, A31571, Life Technologies) with 1:1000 dilution in staining buffer for 2 hours at room temperature and stained with DAPI (300 nM, D1306, Thermo Fisher Scientific) for 10 minutes at room temperature. After washed three times in PBS for 30 min, Cover glasses were mounted onto microscope slides with HydroMount medium (National Diagnostics). Confocal images were acquired using a SP8 confocal microscope (Leica) with a 40x oil objective.

### Quantification of colocalization

The biotinylated monoclonal mouse anti-Syt1 antibody (105 311BT, Synaptic Systems) or the biotinylated anti-VGAT antibody (131 103CpH, Synaptic Systems) was conjugated to streptavidin-conjugated quantum dots (cat. A10196, Thermo Fisher Scientific), and vesicles were loaded as described previously (23). The rabbit polyclonal anti-VGAT antibody conjugated to CypHer5E (cat. 131 103CpH, Synaptic Systems) were used to label inhibitory presynaptic terminals by incubation for 3 hours. QD-Syt1‒ loaded vesicles localized to VGAT-CypHer5E‒labeled presynaptic boutons were manually counted based on colocalization between the QD and CypHer5E fluorescence signals.

## Supporting information

This article contains supporting information available online.

## Acknowledgments

We thank Ching Yeung Fan, Cheuk Long Frank, Lee, and Keun Yang Park for help with data analysis, and members of Park lab for helpful discussion and comments. We also thank Chenglu Tao for discussion. We thank Dr. Curtis Barrett for critically reading the manuscript and providing constructive comments. This work was supported by the Research Grants Council of Hong Kong (Grants 26101117, 16101518, A-HKUST603/17, and N_HKUST613/17) and the Innovation and Technology Commission (ITCPD/17-9).

## Declaration of interest

The authors declare no conflict of interest.

## Notes

### Competing Interest Statement

The authors have declared no competing interest.

