## Supplementary Information for "Inhibitory synaptic vesicles have unique dynamics and exocytosis properties"

**This PDF file includes:**

Figures S1 to S4

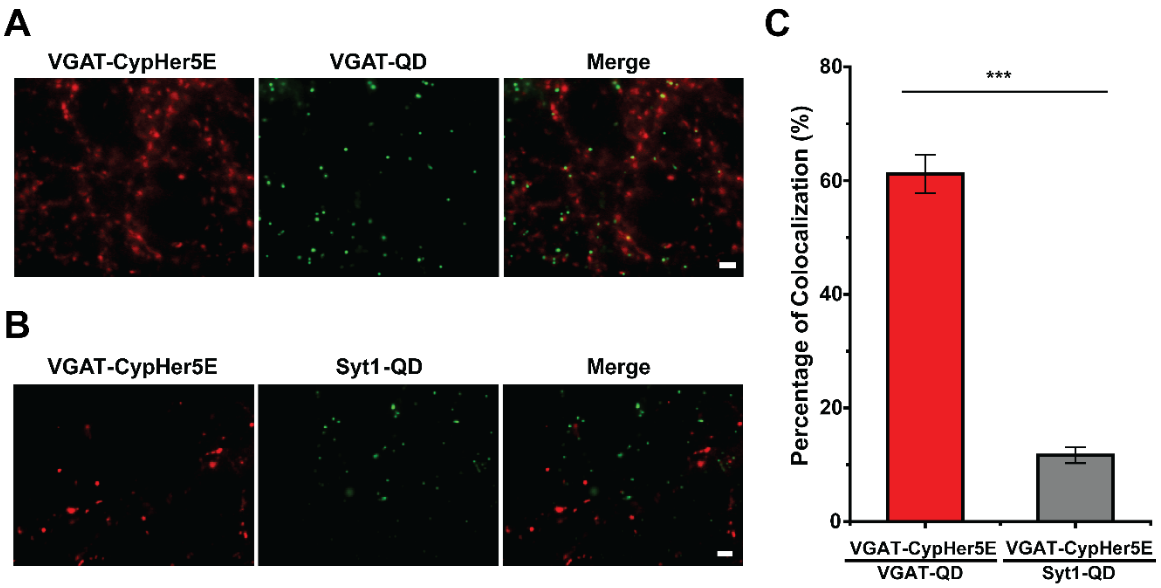


**Fig. S1. VGAT-conjugated QDs preferentially label inhibitory synaptic vesicles in cultured hippocampal neurons. (A-B)** Representative fluorescence images of inhibitory presynaptic terminals labeled with CypHer5E-conjugated antibodies against the luminal domain of VGAT (VGAT-CypHer5E, red) and synaptic vesicles loaded with QDs conjugated to antibodies against the luminal domain of VGAT (VGAT-QD, green; A) or antibodies against Syt1 (Syt1-QD, green; B). Scale bars: 3 µm. **(C)** Percentage of colocalization between VGAT-CypHer5E–labeled boutons and either VGAT-QD–loaded vesicles (61 ± 3.4 % (N = 13 images)) or Syt1-QD–loaded vesicles (12 ± 1.4 % (N = 13)). ***$p$<0.001 (Student's $t$-test).

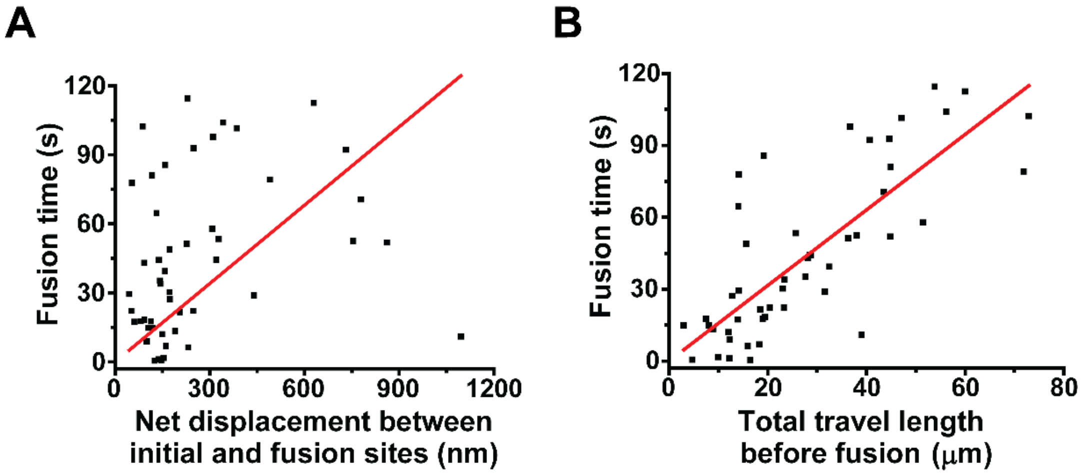


**Fig. S2. Fusion time is correlated with both net displacement and total length traveled for Syt1-QD–loaded vesicles. (A)** Fusion time is plotted against the net displacement; each symbol represents an individual Syt1-SQ–loaded vesicle, and the solid red line represents the linear regression (Pearson's r = 0.71; slope = 0.11 ± 0.02 (n = 49 vesicles)). **(B)** Fusion time is plotted against the total travel length; each symbol represents an individual Syt1-SQ–loaded vesicle, and the solid red line represents the linear regression (Pearson's r = 0.92; slope = 1.58 ± 0.09).

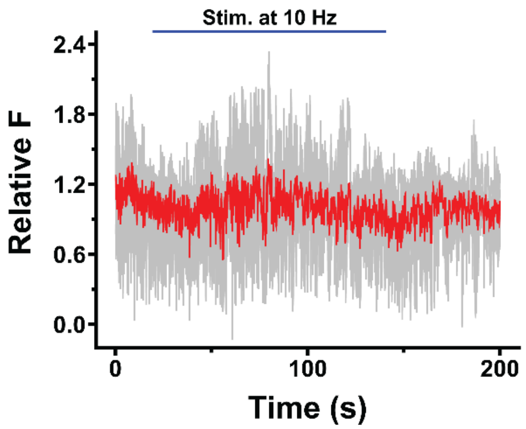


**Fig. S3. Fluorescence traces of non-releasing VGAT-QD–loaded vesicles.** Five individual traces are shown in gray, and the average trace is shown with a red line. A blue horizontal bar represents electrical stimuli (10 Hz) for 120 s starting at 20 s.

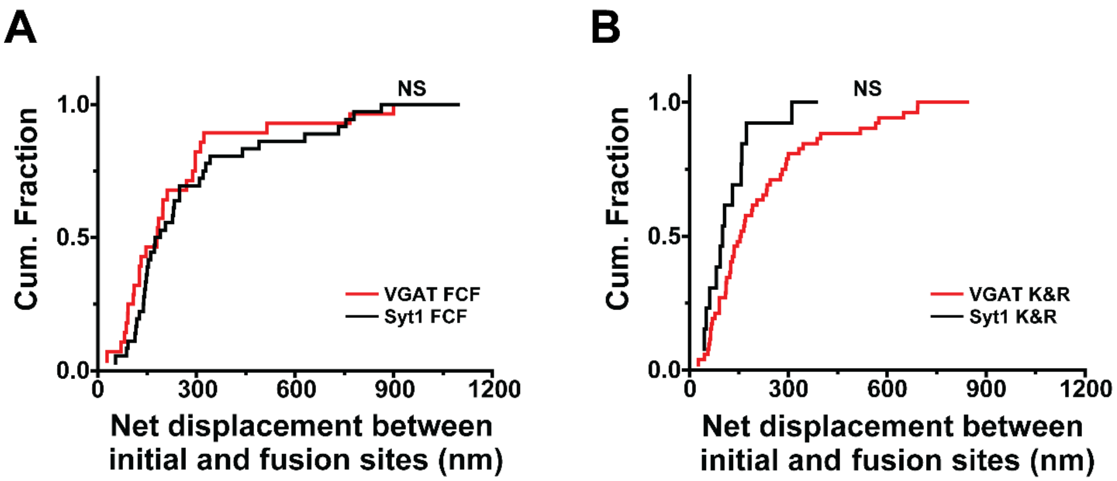


**Fig. S4. Cumulative distribution of net displacement of VGAT-QD–loaded and Syt1-QD–loaded synaptic vesicles undergoing either full-collapse fusion (FCF) or kiss-and-run (K&R) fusion. (A)** Cumulative distribution of net displacement of VGAT-QD–loaded vesicles (n = 28) and Syt1-QD–loaded vesicles (n = 36) undergoing FCF. The net displacement of VGAT-labeled synaptic vesicles undergoing full-collapse fusion (VGAT FCF) was not significantly different from that of Syt1-labeled vesicles undergoing full-collapse fusion (Syt1 FCF) (260 ± 46 nm (n = 28 vesicles) vs. 302 ± 43 nm (n = 36), respectively; $p > 0.4$, K-S test). **(B)** Cumulative distribution of net displacement of VGAT-QD–loaded vesicles (n = 52) and Syt1-QD–loaded vesicles (n = 13) undergoing kiss-and-run (K&R) fusion. The net displacement of VGAT-labeled synaptic vesicles undergoing kiss-and-run fusion (VGAT K&R) was not significantly different from that of Syt1-labeled vesicles undergoing kiss-and-run fusion (Syt1 K&R) (231 ± 27 nm (n = 52) vs. 143 ± 28 nm (n = 13), respectively; $p > 0.3$, K-S test). NS: not significant.
